# Steady at the wheel: conservative sex and the benefits of bacterial transformation

**DOI:** 10.1101/062562

**Authors:** Ole Herman Ambur, Jan Engelstädter, Pål J. Johnsen, Eric L. Miller, Daniel E. Rozen

## Abstract

Many bacteria are highly sexual, but the reasons for their promiscuity remain obscure. Did bacterial sex evolve to maximize diversity and facilitate adaptation in a changing world, or does it instead help to retain the bacterial functions that work right now? In other words, is bacterial sex innovative or conservative? Our aim in this review is to integrate experimental, bioinformatic and theoretical studies to critically evaluate these alternatives, with a main focus on natural genetic transformation, the bacterial equivalent of eukaryotic sexual reproduction. First, we provide a general overview of several hypotheses that have been put forward to explain the evolution of transformation. Next, we synthesize a large body of evidence highlighting the numerous passive and active barriers to transformation that have evolved to protect bacteria from foreign DNA, thereby increasing the likelihood that transformation takes place among clonemates. Our critical review of the existing literature provides support for the view that bacterial transformation is maintained as a means of genomic conservation that provides direct benefits to both individual bacterial cells and to transformable bacterial populations. We examine the generality of this view across bacteria and contrast this explanation with the different evolutionary roles proposed to maintain sex in eukaryotes.

## Introduction

Bacteria have long been appreciated as genomic shape-shifters that gain and lose genes with great regularity [1]. This fluidity is intuitively encapsulated by the idea of the core genome, or the fraction of genes shared by all or most strains of a given species. By this measure only around 50% of the genomic content of many bacterial strains is shared by other strains of the same species, while the rest, the accessory genome, is either unique to a particular genome or shared sporadically across the species [2]. In other estimates, and depending on the species, up to 25% of the genome is the result of horizontal gene transfer (HGT) [3–5]. Although there are wide margins of error on these estimates, owing to difficulties of discerning recent from distant gene transfer events or HGT occurring within or across species, it is clear that HGT plays a key role in shaping bacterial genomes. Equally, the genes subject to HGT seem to have an outsized role on bacterial ecology and evolution, influencing where bacteria are found, what they can consume or degrade, their susceptibility to antibiotics and their virulence as pathogens, among others [3].

Because of these conspicuous benefits, it is easy to draw the conclusion that HGT is uniformly positive, and that bacteria engage in promiscuous sex in order to enhance their adaptability, much like meiotic sex enhances eukaryotic adaptation. However, this conclusion is too narrow and potentially misguided. First, it is important to bear in mind that there is a perception bias with respect to the benefits of HGT because the only instances of HGT that are observed in bacterial genomes are those that have passed the filter of natural selection [6]. Second, just like in eukaryotes, recombination in prokaryotes is associated with several potential costs that limit when, where and how recombination can occur [7,8]. Finally, while there are superficial similarities between eukaryotic sex and recombination in bacteria, the mechanisms underlying these processes are vastly diverged, as are their potential costs and benefits [6]. Thus, although it is tempting to look to eukaryotes to help understand the benefits of sex in prokaryotes, this should be done with caution. The regulation and mechanisms underlying bacterial sex appear to have evolved independently across prokaryotic groups, although some of the core genes associated with recombination are broadly conserved. Also, the frequency of recombination can be highly variable even within a species [6,9,10]. In spite of this, we believe a unifying benefit to bacterial sex can still be found. Here we argue that the benefits of bacterial sex, and transformation in particular, lie in its genomic conservatism and not in the opportunities transformation can provide for genomic innovation. So rather than the “weird sex” observed in some of the other species considered in this Special Issue, we argue that bacteria use “safe sex” to repair DNA or purge deleterious mutations, to overcome stress, and to combat genomic parasites.

Bacteria utilize three primary mechanisms to acquire exogenous DNA as a substrate for recombination: *conjugation*, which is plasmid mediated; *transduction*, which is caused by bacterial viruses called phages; and *transformation*, the regulated uptake and incorporation via homologous recombination of exogenous DNA. Although all three of these mechanisms of recombination contribute to HGT, only natural transformation is exclusively encoded by genes present on the bacterial chromosome. For that reason, transformation is the only process that may have evolved as a form of bacterial sex [6,11]. Accordingly, and as opposed to genes for conjugation and transduction that are predominantly carried by accessory infectious elements, the costs and benefits of transformation are borne solely and directly by the competent bacteria themselves. In addition to the three classic mechanisms of DNA exchange, it has become clear in recent years that DNA may also be transferred between bacteria by other mechanisms, including nanotubes [12], micro-vesicles [13], or gene-transfer agents [14]. However, the prevalence and impact of these agents still remains to be established and we therefore focus on transformation for the remainder of this review.

We first discuss the diverse evolutionary costs and benefits of natural competence. Next, we outline the numerous strategies bacteria use to increase the likelihood that transformed DNA is derived from within the same species. Finally, we consider experimental and theoretical support for the idea of conservative sex and conclude with suggestions for further study. In addition to the discussion below, we also refer interested readers to several excellent reviews that provide more mechanistic or species-specific details of competence induction and transformation [15–17].

## Costs and benefits of natural transformation

### Physiological costs of transformation

Natural transformation is coordinated by a large and complex molecular machinery dedicated to the uptake of DNA from the environment, its intracellular processing and, potentially, its genetic incorporation through recombination. Although there are similarities in the mechanisms of DNA binding, uptake, and incorporation across species, distinct patterns of competence regulation together with the sporadic distribution of competence across bacteria imply that transformation has had multiple independent origins [18,19]. Equally, the rates of natural transformation can vary markedly within a single species [9,10], suggesting that transformation is evolutionarily labile and that bacteria face a trade-off between its costs and benefits. Potential costs are numerous and diverse. The very act of transformation is energetically costly, sometimes involving the transcription of more than 100 genes, only a fraction of which are required for recombination [18]. Moreover, DNA binding, uptake, and incorporation may require a “handling” time that reduces rates of vegetative growth; indeed, in some species competent cells temporarily arrest growth, thereby reducing fitness when in competition with non-competent cells [20]. This cost of entering this persister-like state was proposed to explain why only a fraction of cells in *B. subtilis* become competent [21]. On the other hand, under conditions where rapidly dividing bacteria experience high levels of mortality (e.g., because of antibiotics that target cell wall biosynthesis), the persister-state induced by competence may actually be favoured and thus indirectly select for the maintenance of competence [20]. Another cost arises through recombination itself because recombination is by default a DNA damaging process; evidence suggests that chromosomal integration of DNA that is successfully taken up by the cell can ultimately lead to double stranded breaks that are lethal unless repaired [22,23]. Finally, there are the potentially considerable costs associated with competence-induced cell lysis in e.g. *Streptococcus pneumoniae*, where competent cells actively lyse non-competent members of the same population [24]. Although this last cost is undoubtedly harmful to non-competent cells, it remains possible that competent “killers” benefit by liberating DNA from competing cells, which then serves as a substrate for transformation.

### DNA as food

DNA is metabolically costly and one of the simplest and most intuitive benefits for transformation lies in the possibility that bacteria could use DNA as a resource in the form of nucleotides and nucleotide precursors [25]. Were cells induced to become competent by starvation, DNA could in principle provide sufficient energy for continued replication or repair. Consistent with this possibility, some species, like *Haemophilus influenzae*, require nutritional down-shifts to induce competence and purine depletion activates the competence activator *sxy* [26]. However, several factors argue against this hypothesis as a general explanation for the maintenance of transformation: 1) as yet, there is no clear evidence that the integration of nucleotides taken up by transformation become routed into DNA metabolism; 2) the presence of exogenous DNA does not appear to induce competence in any transformable species; 3) competence in streptococci, like *S. pneumoniae*, is induced for only a short time period during exponential growth when other resources are highly abundant [15]; 4) transported DNA is heavily protected against nuclease digestion within the cell, potentially enabling transported fragments to remain intact as a substrate for recombination [27]; and 5) the hypothesis does not explain why several species that become competent for natural transformation only take up DNA from close relatives due to conserved DNA uptake sequences (DUS) despite the fact that non-homologous DNA could be used as a source of nucleotides for direct use or degradation [19,28]. In addition to these concerns, it remains uncertain, on energetic grounds, if the costs of DNA transport are sufficiently offset by any metabolic savings provided by exogenous DNA [11]. Thus, despite the intuitive appeal of this idea, the evidence in its favour is currently limited.

### DNA repair

An old idea for a potential benefit of transformation is that acquired DNA is used as a substrate for genome repair [29,30]. Early experimental evidence indicated an immediate benefit of DNA uptake on transformant survival relative to the remainder of the population in *Bacillus subtilis* [30–32]. However these earlier results were countered in the same species by evidence that genotoxic stress did not induce competence [33], as predicted by the original idea. A more recent extension of this “DNA repair hypothesis” proposes that transformation is a general stress response [34]. This idea is supported by the fact that some, but not all [35], naturally transformable human pathogens such as *Streptococcus pneumoniae, Legionella pneumophila* and *Helicobacter pylori* lack a general SOS response and that DNA damaging agents, including some antibiotics, induce competence in these species [34,36,37]. Current evidence suggests, however, that the benefits of competence induction in these species may be unlinked from the effects of transformation per se (i.e. DNA uptake and integration) [38,39]. Thus although these reports reveal that at least some forms of stress can induce competence, they do not always provide evidence that this response is adaptive, nor provide clear indications of the mechanisms underlying these benefits. Furthermore, they do not take into account the abundant evidence showing that competence in many species is induced by quorum sensing, irrespective of exogenous stress, and that other unambiguous forms of stress, like temperature and pH, can even repress natural transformation [40–43]. Taken together, the available data do not clearly support the classical “DNA for repair” hypothesis; instead they are more consistent with the more general hypothesis that competence development in itself is beneficial, albeit under restricted conditions.

### Homologous recombination

Natural transformation has the potential to shuffle alleles at different loci within a population. This gives rise to a suite of potential benefits and costs of transformation associated with genetic recombination that are largely identical to those studied for meiotic sex in eukaryotes. Below, we briefly review two of the principles – epistasis and Hill-Robertson interference – through which transformation may be favoured, in both cases through either helping to purge deleterious or fix beneficial mutations. For more in-depth reviews of this large field we refer to Otto [44], Hartfield & Keightley [45], and, in the context of bacteria, Vos [46].

Selection with epistasis, i.e. non-independent fitness effects of alleles at different loci, produces nonindependent gene associations within a population (linkage disequilibria). At the most basic level, recombination can have a detrimental effect as it breaks up co-adapted gene complexes (e.g. [47]). Nevertheless, if deleterious mutations that interact with negative epistasis continually arise within a population, recombination can be favoured because it can increase genetic variance and therefore the efficacy by which natural selection purges deleterious mutations (the ‘deterministic mutation hypothesis’) [48]. This was shown theoretically to also provide a benefit to natural transformation in bacterial populations [49,50]. However, empirical work has shown that while negative epistasis sometimes exists, it is far from pervasive and there are many systems in which positive or no epistasis was reported (reviewed in [51,52]). For this reason, the deterministic mutation hypothesis has been broadly disregarded as a main contender to explain the ubiquity of sex, including natural transformation.

A second class of explanations for why recombination can be beneficial relies on the Hill-Robertson effect [53,54]. Here, an interaction between natural selection and stochastic effects in finite populations (through random genetic drift or mutation) produces genetic associations (negative linkage disequilibria) that reduce genetic variance for fitness. By breaking up these associations, recombination can increase the efficacy of natural selection and genes that increase the recombination rate can be indirectly selected for. Advantages of recombination stemming from the Hill-Robertson effect come in different forms, and may involve both beneficial and deleterious mutations (e.g., [55,56]). Extreme manifestations are the Fisher-Muller model [57,58], in which recombination brings together beneficial mutations that would in asexual populations compete with each other (‘clonal interference’), and Muller’s ratchet [59], the perpetual loss of mutation-free individuals in asexual populations subject to deleterious mutations. A number of mathematical and simulation models have been developed to specifically investigate variants of the Hill-Robertson effect in the context of bacterial sex, confirming that natural transformation can be favoured both in populations subject to recurring deleterious mutations [21,60] and in adapting bacterial populations [21,61,62].

Experimental work testing these ideas has focused on the Fisher-Muller model, the central prediction of which is that recombination accelerates the adaptation rate of sexual relative to asexual variants [57,58]. Evolution experiments with the Gram-negative bacterium *Escherichia coli* [63] undergoing plasmid-mediated recombination and the yeast *Saccharomyces cerevisiae* [64] support this hypothesis, while both reports strongly suggest the crucial role of recombination in relieving the effects of clonal interference. Reduced clonal interference has also been proposed as a key genetic mechanism underlying the evolutionary maintenance of natural transformation [46]. For example, transformable *Helicobacter pylori* adapted more rapidly than non-transformable, otherwise isogenic, populations [65]. By contrast, a study using the highly transformable species *Acinetobacter baylyi* failed to detect any consistent evolutionary advantage of natural transformation during 1000 generations of experimental evolution [66]. It was later demonstrated that the benefits of natural transformation in *A. baylyi* are growth-phase dependent, whereby a positive effect was demonstrated during active growth/early stationary phase which was offset by reduced adaptation to the experimental conditions during stationary/death phase [67]. Similar context-dependent benefits of transformation were shown in the human pathogen *Streptococcus pneumoniae*. Here, competence for natural transformation was disadvantageous when populations evolved in benign experimental conditions, but not when they evolved in the presence of periodic mild stress (sub-inhibitory concentrations of kanamycin) [68]. In addition, evolving competent *S. pneumoniae* populations fixed significantly fewer mutations than isogenic non-competent lineages, while non-competent lineages were more likely to evolve mutator genotypes [68]. Taken together, the few experimental tests of Fisher-Muller advantages to bacterial transformation provide a mixed picture: while some studies support the idea that natural transformation can accelerate adaptation, other results highlight that these responses are either context dependent or absent altogether (at least under the conditions examined).

### Biased allelic replacement

Because of the symmetry of meiotic crossovers, the central influence of eukaryotic recombination at the population level is to reduce linkage disequilibria. By contrast, natural transformation can also affect allele frequencies within a population because of its inherent asymmetry: genetic material is taken from a donor gene pool of free DNA molecules in the environment (“eDNA”) and replaces material within a recipient gene pool of living cells, and those two gene pools are not necessarily identically in terms of allelic composition. This was first recognized by Redfield [69] who showed with a mathematical model that if bacteria carrying deleterious alleles are more likely to die and thereby release their DNA into the environment than wild-type bacteria, this creates a bias towards taking up deleterious alleles and thus a distinctive cost of natural transformation that is independent from any effects derived from gene shuffling. More recently, models of well-mixed and spatially structured populations were developed that explicitly incorporated a pool of free eDNA subject to decay [62,70]. These models recovered a similar detrimental effect of transformation in populations adapting to new environmental conditions because the eDNA may build up an over-representation of old, non-beneficial alleles. Interestingly, however, taking up and incorporating “old alleles” may also turn out to be beneficial in situations where environmental conditions change frequently. According to this idea of “genetic time travel”, natural transformation may allow the bacteria to make use of a reservoir of genetic material preserved in the form of eDNA [71].

### Defence against genomic parasites

Recently, yet another hypothesis for the raison d’être of natural transformation has been proposed [72], according to which transformation helps bacteria to cure their genomes of costly integrated mobile genetic elements (MGEs) such as phages or conjugative elements. This hypothesis is based on the observation that successful transformation only requires homology between short stretches of DNA at the ends of the transformed fragment and the host chromosome, while the regions in the middle can freely vary [73,74]. As a consequence, transformation can lead to either the incorporation of new genetic material (as is usually emphasized) or its expulsion, if, for example the transforming DNA lacks a MGE that is present in the transformed recipient. But under what conditions would this work? Croucher *et al.* [72] motivate their hypothesis with the observation that in both *S. pneumoniae* and *H. influenzae*, the length of newly incorporated DNA fragments approximately follows a geometric distribution [75,76]. They argue that as a consequence of this bias transformed cells will be more likely to incorporate short rather than long DNA stretches, in turn preferentially leading to the loss of MGE. However, generalizing their idea, we suspect that such a bias may not be strictly necessary. Although transformation with long DNA fragments can lead to MGE acquisition, it can also – and potentially at the same rate – lead to their loss if transformed fragments cover the insertion site of the MGE. Conversely, transformation with DNA fragments shorter than the MGE can only ever lead to MGE loss. Thus, as long as there is any natural transformation with DNA fragments that are shorter than the MGE in question, there should be an automatic bias towards exclusion of the element. The existence of this bias should then be independent of the size distribution of incorporated DNA, even though its magnitude may still be affected by the size distribution. This more general version of the defence hypothesis remains to be specifically tested.

In addition to corroborating their hypothesis by means of a mathematical model, Croucher *et al.* [72], also showed through whole-genome analyses that transformable pneumococci harboured fewer prophages than non-transformable ones and that phages often insert into genes involved in the DNA uptake machinery, thus disrupting the putative cellular defence mechanism against MGE. In a small preliminary analysis for this review, we screened the whole genomes of 10 strains of *S. pneumoniae* for insertion sequences, transposons and phages (putative and confirmed) [77] from which we have precisely quantified transformation rates [10]; while 5 of the strains are unable to become competent, the other 5 show variable rates of transformation across several orders of magnitude. Consistent with the predictions of the Croucher *et al.* model we observed clear but non-significant negative correlations between transformation rate and the numbers of putative and confirmed phage. Although no strong conclusions can be drawn from these limited tests, we believe this approach is likely to be very powerful for further tests of these ideas, as it can tie experimentally validated differences in transformation rates to the number of mobile elements carried in a given genome.

### The potential for consensus

Despite the numerous studies of competence across a broad swath of competent bacterial species, there still remains considerable uncertainty about its function because no single explanation seems to capture the particulars of different species growing in different contexts. And this may in fact be the answer: as with eukaryotic sex, a plurality of explanations must be considered [78]. While Haemophilus may become competent when it is hungry or to repair DNA damage, streptococci, *H. pylori* [36], and *Legionella pneumophiliae* [37,79] may only do so when they are damaged, while *Acinetobacter baylyi* [42] and *Neisseria meningitidis* [80] are constitutively competent (although not necessarily all cells in the population at the same time). However, a consensus option may lie in the fact that not all transformable DNA is of equal appeal or value to each bacterial species. Most obviously, non-homologous DNA is a poorer substrate for recombination than homologous DNA. But it goes further than this. As we detail below, all competent bacterial species go to extreme lengths to ensure that the DNA they take up or recombine is either from the same species or from the same strain. This source of DNA is a safe and conservative way to either repair DNA or purge deleterious mutations (including mutator alleles [68]) by restoring the wild-type allele, to re-establish functions from related strains that have already been tested in a highly similar genetic background or to purge genomes of potentially harmful parasitic elements. This conservative explanation builds on the multilayered barriers in place to reduce outcrossing, and offers a general and testable hypothesis for the role of transformation in bacteria.

### Barriers to transformation outside and within the cell

Barriers to transformation are mediated by passive and active processes that serve to dramatically increase the likelihood that competent cells will be transformed with DNA from clonemates. While passive barriers offer approximate routes to restrict transformation, they are imprecise. By contrast, active restrictions to DNA recognition, uptake and recombination are significantly more limiting, and together these redundant processes strongly bias which DNA becomes available for transformation, relative to the broader pool of eDNA.

#### Barriers acting outside the cell

Ecological barriers to recombination require no special mechanisms of exclusion and arise as a simple consequence of the fact that persistence of eDNA is limited and because bacterial growth is spatially structured. For these passive reasons, HGT will tend to take place between cells that are in close proximity to one another and thus often clonal. At a broad scale, this leads to networks of gene exchange that are ecologically confined. For example, by examining routes of HGT via a network approach, Popa and Dagan [7] estimated that 74% of identified HGT events occurred among bacteria residing in the same habitat. Similarly, Smillie *et al.* [81] found that bacteria in the human microbiome residing in the same body sites were more likely to exchange genes than those from different sites. Although compelling, these examples deal with HGT broadly, and may only be partly due to transformation. However, even at this finer scale, proximity, together with other processes examined below, is likely to increase the probability of clonal transfer. *Vibrio cholerae*, for example, becomes competent in the presence of chitin, a substance these bacteria encounter during growth on the surface of zooplankton [82]. Adding to this, Vibrio competence is induced by a quorum-sensing system that is activated by high concentrations of a secreted chemical signal that serves as a proxy measure for high densities of cells of the same species that share this signal [83].

Several other naturally competent species rely on similar modes of species or clone-level recognition to regulate induction. The Gram-positive species *B. subtilis* and *S. pneumoniae* become competent following detection of a threshold concentration of a secreted peptide signal that is species specific [16,34]. Because the signal is produced constitutively, its environmental concentration approximately scales with the density of producing cells [15,84]. While quorum sensing is generally sufficient to ensure induction is coordinated among cells of the same species, it would not allow finer coordination within species. A possible solution to this problem lies in the fact that *B. subtilis* and *S. pneumoniae* are polymorphic for the peptide signals they produce (called pherotypes) [85,86], and there is no apparent cross-talk between a given peptide and non-cognate receptors [87]. In *S. pneumoniae*, several groups have argued that this type of polymorphism could permit transformation to occur preferentially among cells expressing the same pherotype—much like bacterial mating types, although current results on this issue are mixed [85,88].

By using ecological proximity and quorum sensing systems to estimate cell density, bacteria can regulate competence so that it is only turned on when cells are surrounded by individuals of the same species. More strikingly, some naturally competent bacterial species can also strongly influence which DNA becomes available as substrate for transformation by co-ordinately regulating competence induction with diverse modes of cell killing. Some streptococci, for example, use a process dubbed “fratricide” whereby a sub-population of competent cells kill and lyse non-competent members of the same population [24, 89]. In *S. pneumoniae*, competence coincides with the production of both hydrolytic enzymes that degrade the cell wall and also the secretion of multiple classes of bacteriocins, including the highly diverse *blp* (bacteriocin-like peptide) gene-cluster [90–93]. Importantly, killer cells are protected from the action of these secreted weapons, which leads to a one-way routing of DNA from target cells to transformed recipients. Another recent discovery confirms this coincident expression of competence and killing, but via an entirely distinct mechanism. Many Gram-negative bacteria express a needle-like machine called a Type IV secretion system (T6SS) that injects toxins into non-immune target cells, resulting in their death and lysis [94]. Remarkably, *V. cholera* co-regulates competence induction with T6SS-mediated predation after which the liberated DNA is immediately available for uptake and incorporation [95]. As with fratricide, this form of predation is not a fail-safe means by which cells can acquire clonal DNA; however, together with ecological proximity and quorum sensing-dependent regulation of competence, clonal uptake is likely to be the most common outcome.

While external barriers to transformation may be reliable, they are also susceptible to failure in multispecies bacterial communities where eDNA will inevitably be present due to the continuous lysis of dead cells. To offset these risks, bacteria have evolved a suite of redundant mechanisms that can limit transformation by excluding cells on the basis of specific sequence tags, called uptake sequences, or by restricting incorporation or expression following DNA update.

#### Sequence tags restricting DNA uptake

Several species in two phylogenetically distinct bacterial families, the Pasteurellaceae and the Neisseriaceae, have evolved sequence specific uptake of DNA [19,28,96]. In short, these sequences act like small barcodes that identify that the DNA a competent cell binds comes from the correct species. Discrimination between “self” and “other” DNA occurs at the bacterial surface. In the Neisseriaceae, the small competence protein ComP binds a small signature in DNA and initiates uptake [97]. A similar factor has not yet been identified in the Pasteurellaceae and the exact molecular interactions responsible for sequence specificity awaits characterization in this family.

Nonetheless, the genomes of these competent bacteria carry hundreds to several thousands of these small specific signatures in DNA termed DNA Uptake Sequences (DUS) in the Neisseriaceae [98] and Uptake Signal Sequences (USS) in the Pasteurellaceae [28]. Typically, DUS extend for 12 nucleotides with the following consensus sequences: 5’-ATGCCGTCTGAA-3’ and USS 9 nt 5’-AAGTGCGGT-3’, with a defined core of 3-4 essential nucleotides, although different dialects of both DUS and USS exist in different species [99,100]. USS and DUS are not homologous, suggesting that the uptake specificities in the Neisseriaceae and the Pasteurellaceae are the results of convergent evolution. The DUS/USS are spread throughout their respective chromosomes so that most parts of the genomes are likely substrates for transformation. There is a close concordance between DUS distribution and conversion fragments identified from whole genome alignments [101]. In some species the number of DUS are so high that they occupy more than 3% of the entire genome, reflecting the substantial genetic investment in maintaining specific transformation [99]. How high is the DUS/USS barrier against non-DUS/USS DNA? The dependency for DUS in transformation has been shown to vary from absolute to less pronounced between strains [96]. The location of DUS relative to homologous and heterologous stretches of DNA, together with DUS/USS integrity, and strandedness also influence transformation efficacy [96,99,102–104]. However, DUS/USS always biases transformation to involve homologous DNA. DUS/USS are remarkably conserved and single deviations from the signal are rare in their respective genomes and experimentally reduce or eliminate transformability [99,100]. Interestingly, DUS are clustered in the core genome [101] and both DUS and USS are biased towards genes associated with genome maintenance [105] Since these important and often essential genes of the core genome carry evolved and costly signatures for transformation, often inside coding regions, an association between their conserved status and transformation may therefore exist. By contrast, DUS are underrepresented in genes encoding surface exposed epitopes that typically are hypervariable and in the accessory genome. DUS and transformation in these species are therefore not apparently associated with variability (e.g. for immune evasion) and supports a conservative rationale for transformation.

#### Restriction modification systems (RMS)

Once in the cytoplasm, and in species that do not rely on DUS/USS for discrimination, DNA faces another threat to its integrity: restriction modification systems (RMS). Typically RMS are constituted by a restriction endonuclease that cuts double-stranded DNA (dsDNA) at non-methylated specific recognition sites and a corresponding methylase that can protect the same recognition site. RMS are nearly ubiquitous in bacteria [106] where they raise barriers against the stable integration of alien DNA from transduction, conjugation and transformation [8]. More than 5000 RMS are biochemically or genetically characterized and even more are added to the comprehensive RMS database REBASE daily [107]. At which exact stage during transformation the restriction endonuclease cuts DNA has been the matter of some debate because the endonucleolytic activity is clearly directed towards dsDNA, whereas it is ssDNA that enters the cytoplasm according to current models for transformation [16]. A postreplication and hence postrecombination model has been proposed where transformed chromosomes are subject to endonucleolytic attack once double-strandedness has been restored by chromosomal replication [74]. Using commercially available methylases it has become possible to specifically study the extent of individual barriers to transformation due to RMS in many different bacteria such as *Pseudomonas stutzeri* (e.g. [108]). Specific knock-outs of restriction endonucleases or specific pre-methylation of transforming DNA has also revealed several further species that can become competent for transformation, such as the human pathogen *Staphylococcus aureus* [109,110] and the thermophilic cellulolytic *Caldisiruptor bescii* [111]. In this latter organism, the restriction barrier is apparently absolute when using transforming homologous DNA propagated in another species (*E. coli*). Many other bacteria display very high and some absolute RMS-barriers to transformation that depends on the DNA source, sequence and homology [102,112]. As discussed above, competence may be regulated and induced in some bacteria when conditions for transformation are favorable. About half of the sequenced *S. pneumoniae* strains harbour an unusual ssDNA methylase, DpnA that protects incoming DNA from restriction at 5’-GATC-3’ sites by DpnII during transformation. Notably, DpnA forms part of the competence regulon and is expressed from an competence-inducible promotor [27,74]. *S. pneumoniae* can thereby transiently lower the transformation barrier in periods when competence is induced.

By limiting stable integration of heterologous DNA and favouring homologous DNA of equal modification status, RMS may therefore be important drivers of speciation in bacteria [113]. Genomic studies of the naturally competent *N. meningitidis* has indeed shown that divergent RMS profiles in different lineages of this species have contributed to the sexual isolation of phylogenetic clades [114]. In addition to RMS, the bacterial immune system based on clustered, regularly interspaced, short palindromic repeats (CRISPRs) found in 40% of all bacteria [115] have been shown to prevent natural transformation in a streptococcal transformation/infection model [116]. A genomics study of the naturally competent *Aggregatibacter actinomycetemcomitans* showed that the loss of competence was followed by the loss of CRISPRs in an evolutionary time frame [117] linking the two phenomena. However, the evolutionary consequence of CRISPRs inhibiting horizontal gene transfer remains a matter of debate [118].

#### Mismatch repair

Homologous recombination (HR) is the last step in transformation where incoming DNA is exchanged with a similar or identical allele in the chromosome. A HR event that involves two identical segments of DNA leaves no genetic signature thereby hiding it from retrospective detection. By contrast, HR involving similar but not identical alleles is traceable and has hence been more studied. This in turn may have fostered the common view that transformation evolved to innovate and not to conserve. HR is ubiquitous in nature and in *E. coli* more than 25 genes are involved in the process [119]. During transformation incoming ssDNA is coated with single stranded binding protein (SSB) and sequentially the recombination mediator DprA [120] or RecO facilitates loading of the recombinase RecA that ultimately leads to strand exchange [121,122]. RecA allows the ssDNA to match up with a homologous segment in the chromosome with high fidelity and to initiate the allelic exchange [123]. RecA has been shown able to very rapidly identify its relatively small homologous target in a suspension where heterologous DNA targets are in a 200,000-fold excess [124]. Homology must not be complete for strand exchange to take place and variable degrees of RecA fidelity have been documented. However, a strong and persistent negative exponential correlation between sequence divergence and transformation has been shown in different competent bacteria and archaea [125–128]. Furthermore, the recombination potential between particular loci of divergent, but related, Acinetobacter species has been shown to be dependent on the composition of DNA sequences up to several kilobases away from the loci in consideration [129]. Also, mismatch repair (MMR) systems may control the tolerance for heterology in HR. In the competent *B.subtilis, S. pneumoniae* and *P. stutzeri* MMR plays a lesser role in controlling mismatches than in the non-or less-competent species such as *E. coli*. In these competent species RecA is the main discriminator against sequence heterology during HR [73,130].

#### HN-S associated gene silencing

Although the barriers described above are efficient, successful transfers of DNA between species and genera regularly occur and are important drivers of microbial evolution. DNA can be mobilized by means of plasmids, transposons and bacteriophages [131]. Regions in bacterial chromosomes of foreign and phylogenetically distant (xenogeneic) origin can be detected by searching for particular signatures such as unusual base composition (GC-content). The base composition in such regions ameliorate over time and “tune” into the genomic signature of its new host since they experience the same mutational processes affecting all genes in the recipient genome [3]. Notably, both Gram-positive and Gram-negative bacteria have evolved repressors that target such non-ameliorated xenogeneic DNA [132,133]. In effect, these repressors silence the expression of genes that could cause harmful effects in the host organism. However, the longer-term fate of such silenced regions is not well understood, and it could even be that such gene silencing mitigates costs of carrying novel genes thereby facilitating their longer-term persistence. Alternatively, silenced regions may be more likely to acquire nonsense mutations, as there is little selection for the maintenance of these unexpressed genes.

#### Fitness costs

In the event that passive and active barriers to transformation do not prevent transformation of non-homologous sequences, transformed cells may still fail to persist and spread. Focusing on HGT generally, Baltrus [134] outlined a series of cellular costs associated with recombination of foreign DNA. Among these, new sequences can cause toxicity due to protein misfolding, misregulation or inefficient translation or transcription due to different GC content (if not adequately silenced by HNS) or misregulation. More simply, transformed DNA can potentially displace functional alleles that are highly coevolved with the host genome, a history that will not exist for diverged sequences. Although these diverse fitness costs are difficult to quantify, several studies have examined the issue by transforming recipient cells (not necessarily natural competent ones) with donor DNA of different size and origins. Knoppel *et al.* [135] found that of 90 such transfers, surprisingly few introduced significant costs while most were neutral. A clear caveat of this work, however, is that fitness was measured during short-term assays with a detection limit that may fail to identify costs that are relevant over longer periods. At the same time, over the longer term and like antibiotic resistance, fitness costs of transformation can potentially be compensated by second-site mutations or by amelioration of the transformed sequence itself [136].

### Conclusions and Outlook

Transformation, together with other modes of HGT, can undoubtedly have an enormous influence on bacterial genomes and ecology. It can facilitate invasion into novel habitats and permit escape from drug or immune pressure [3,46]. However, these benefits are probably more the exception than the rule. The vast majority of HGT events between different strains or species of bacteria are expected to be deleterious, and this has given rise to a complex suite of processes that limit transformation to close relatives with highly similar genomes and ecologies. Moreover, even when acting within largely clonal populations, there is currently no consistent evidence that natural transformation enables bacteria to become more fit or adapt more rapidly to their environment. Together, this leads us to suggest that transformation will tend to be a conservative mechanism, acting much like any other repair process in the cell. This expanded idea of repair can extend to specific forms of DNA damage, deleterious mutations, to more generic stress, or to the removal of genomic parasites. However, even if this is the predominant role for transformation, it clearly does not preclude that transformed DNA will also occasionally provide important adaptive benefits at the individual and population levels. But this leads to important unresolved questions: 1) if transformation is beneficial because of its capacity to repair damage (defined most broadly), why is natural competence so sparsely distributed across bacteria; 2) related to this, why are transformation rates so highly variable within naturally competent species; and 3) what factors, if any, unite species that retain natural competence? The challenge for future work is to address these questions by integrating theoretical and bioinformatics approaches, as exemplified by the recent paper from Croucher *et al.* [72]. There is also further need to consider how the three modes of HGT differentially influence the dynamics of the core and accessory genomes of competent species. Finally, we encourage the development of additional long-term experimental studies that address the benefits of competence induction and transformation for stress-resistance and genome maintenance at short and evolutionary time-scales.

## Additional Information

### Authors’ Contributions

All authors contributed equally to the structure and writing of this review and approved of the final submission. EM conducted bioinformatic analyses of pneumococcal phage.

## References

1. Vos, M., Hesselman, M. C., te Beek, T. A., van Passel, M. W. J. & Eyre-Walker, A. 2015 Rates of Lateral Gene Transfer in Prokaryotes: High but Why? Trends Microbiol. 23, 598–605. (doi:10.1016/j.tim.2015.07.006)

2. Segerman, B. 2012 The genetic integrity of bacterial species: the core genome and the accessory genome, two different stories. Front. Cell. Infect. Microbiol. 2, 116. (doi:10.3389/fcimb.2012.00116)

3. Ochman, H., Lawrence, J. G. & Groisman, E. A. 2000 Lateral gene transfer and the nature of bacterial innovation. Nature 405, 299–304. (doi:10.1038/35012500)

4. Lapierre, P. & Gogarten, J. P. 2009 Estimating the size of the bacterial pan-genome. Trends Genet. 25, (107–10). (doi:10.1016/j.tig.2008.12.004)

5. Nakamura, Y., Itoh, T., Matsuda, H. & Gojobori, T. 2004 Biased biological functions of horizontally transferred genes in prokaryotic genomes. Nat. Genet. 36, (760–6). (doi:10.1038/ng1381)

6. Redfield, R. J. 2001 Do bacteria have sex? Nat. Rev. Genet. 2, (634–9). (doi:10.1038/35084593)

7. Popa, O. & Dagan, T. 2011 Trends and barriers to lateral gene transfer in prokaryotes. Curr. Opin. Microbiol. 14, (615–23). (doi:10.1016/j.mib.2011.07.027)

8. Thomas, C. M. & Nielsen, K. M. 2005 Mechanisms of, and barriers to, horizontal gene transfer between bacteria. Nat. Rev. Microbiol. 3, (711–21). (doi:10.1038/nrmicro1234)

9. Maughan, H. & Redfield, R. J. 2009 Extensive variation in natural competence in Haemophilus influenzae. Evolution 63, (1852–66). (doi:10.1111/j.1558-5646.2009.00658.x)

10. Evans, B. A. & Rozen, D. E. 2013 Significant variation in transformation frequency in Streptococcus pneumoniae. ISME J. 7, (791–9). (doi:10.1038/ismej.2012.170)

11. Seitz, P. & Blokesch, M. 2013 Cues and regulatory pathways involved in natural competence and transformation in pathogenic and environmental Gram-negative bacteria. FEMS Microbiol. Rev. 37, (336–63). (doi:10.1111/j.1574-6976.2012.00353.x)

12. Dubey, G. P. & Ben-Yehuda, S. 2011 Intercellular nanotubes mediate bacterial communication. Cell 144, (590–600). (doi:10.1016/j.cell.2011.01.015)

13. Fulsundar, S., Harms, K., Flaten, G. E., Johnsen, P. J., Chopade, B. A. & Nielsen, K. M. 2014 Gene transfer potential of outer membrane vesicles of Acinetobacter baylyi and effects of stress on vesiculation. Appl. Environ. Microbiol. 80, (3469–83). (doi:10.1128/AEM.04248-13)

14. Lang, A. S., Zhaxybayeva, O. & Beatty, J. T. 2012Gene transfer agents: phage-like elements of genetic exchange. Nat. Rev. Microbiol. 10, (472–82). (doi:10.1038/nrmicro2802)

15. Straume, D., Stamsås, G. A. & Håvarstein, L. S. 2015 Natural transformation and genome evolution in Streptococcus pneumoniae. Infect. Genet. Evol. 33, (371–80). (doi:10.1016/j.meegid.2014.10.020)

16. Dubnau, D. 1991 Genetic competence in Bacillus subtilis. Microbiol. Rev. 55, 395–424.

17. Metzger, L. C. & Blokesch, M. 2015 Regulation of competence-mediated horizontal gene transfer in the natural habitat of Vibrio cholerae. Curr. Opin. Microbiol. 30, (1–7). (doi:10.1016/j.mib.2015.10.007)

18. Johnsborg, O., Eldholm, V. & Håvarstein, L. S. 2007 Natural genetic transformation: prevalence, mechanisms and function. Res. Microbiol. 158, (767–78). (doi:10.1016/j.resmic.2007.09.004)

19. Mell, J. C. & Redfield, R. J. 2014 Natural competence and the evolution of DNA uptake specificity. J. Bacteriol. 196, (1471–83). (doi:10.1128/JB.01293-13)

20. Johnsen, P. J., Dubnau, D. & Levin, B. R. 2009 Episodic selection and the maintenance of competence and natural transformation in Bacillus subtilis. Genetics 181, (1521–33). (doi:10.1534/genetics.108.099523)

21. Wylie, C. S., Trout, A. D., Kessler, D. A. & Levine, H. 2010 Optimal strategy for competence differentiation in bacteria. PLoS Genet. 6, e1001108. (doi:10.1371/journal.pgen.1001108)

22. Kickstein, E., Harms, K. & Wackernagel, W. 2007 Deletions of recBCD or recD influence genetic transformation differently and are lethal together with a recJ deletion in Acinetobacter baylyi. Microbiology 153, (2259–70). (doi:10.1099/mic.0.2007/005256-0)

23. Harms, K., Schön, V., Kickstein, E. & Wackernagel, W. 2007 The RecJ DNase strongly suppresses genomic integration of short but not long foreign DNA fragments by homology-facilitated illegitimate recombination during transformation of Acinetobacter baylyi. Mol. Microbiol. 64, (691–702). (doi:10.1111/j.1365-2958.2007.05692.x)

24. Steinmoen, H., Knutsen, E. & Håvarstein, L. S. 2002 Induction of natural competence in Streptococcus pneumoniae triggers lysis and DNA release from a subfraction of the cell population. Proc. Natl. Acad. Sci. U. S. A. 99, (7681–6). (doi:10.1073/pnas.112464599)

25. Redfield, R. J. 1993 Genes for breakfast: the have-your-cake-and-eat-it-too of bacterial transformation. - PubMed - NCBI. Heredity (Edinb). 84, 400–404.

26. Sinha, S., Mell, J. & Redfield, R. 2013 The availability of purine nucleotides regulates natural competence by controlling translation of the competence activator Sxy. Mol. Microbiol. 88, (1106–19). (doi:10.1111/mmi.12245)

27. Johnston, C., Polard, P. & Claverys, J.-P. 2013 The DpnI/DpnII pneumococcal system, defense against foreign attack without compromising genetic exchange. Mob. Genet. Elements 3, e25582. (doi:10.4161/mge.25582)

28. Danner, D. B., Deich, R. A., Sisco, K. L. & Smith, H. O. 1980 An eleven-base-pair sequence determines the specificity of DNA uptake in Haemophilus transformation. Gene 11, 311–8.

29. Michod, R. E., Bernstein, H. & Nedelcu, A. M. 2008 Adaptive value of sex in microbial pathogens. Infect. Genet. Evol. 8, (267–85). (doi:10.1016/j.meegid.2008.01.002)

30. Michod, R. E., Wojciechowski, M. F. & Hoelzer, M. A. 1988 DNA repair and the evolution of transformation in the bacterium Bacillus subtilis. Genetics 118, 31–9.

31. Hoelzer, M. A. & Michod, R. E. 1991 DNA repair and the evolution of transformation in Bacillus subtilis. III. Sex with damaged DNA. Genetics 128, 215–23.

32. Wojciechowski, M. F., Hoelzer, M. A. & Michod, R. E. 1989 DNA repair and the evolution of transformation in Bacillus subtilis. II. Role of inducible repair. Genetics 121, 411–22.

33. Redfield, R. J. 1993 Evolution of natural transformation: testing the DNA repair hypothesis in Bacillus subtilis and Haemophilus influenzae. Genetics 133, 755–61.

34. Claverys, J.-P., Prudhomme, M. & Martin, B. 2006 Induction of competence regulons as a general response to stress in gram-positive bacteria. Annu. Rev. Microbiol. 60, (451–75). (doi:10.1146/annurev.micro.60.080805.142139)

35. Sweetman, W. A., Moxon, E. R. & Bayliss, C. D. 2005 Induction of the SOS regulon of Haemophilus influenzae does not affect phase variation rates at tetranucleotide or dinucleotide repeats. Microbiology 151, (2751–63). (doi:10.1099/mic.0.27996-0)

36. Dorer, M. S., Fero, J. & Salama, N. R. 2010 DNA damage triggers genetic exchange in Helicobacter pylori. PLoS Pathog. 6, e1001026. (doi:10.1371/journal.ppat.1001026)

37. Charpentier, X., Kay, E., Schneider, D. & Shuman, H. A. 2011 Antibiotics and UV radiation induce competence for natural transformation in Legionella pneumophila. J. Bacteriol. 193, (1114–21). (doi:10.1128/JB.01146-10)

38. Engelmoer, D. J. P. & Rozen, D. E. 2011 Competence increases survival during stress in Streptococcus pneumoniae. Evolution 65, (3475–85). (doi:10.1111/j.1558-5646.2011.01402.x)

39. Mongold, J. A. 1992 DNA repair and the evolution of transformation in Haemophilus influenzae. Genetics 132, 893–8.

40. Chen, J. D. & Morrison, D. A. 1987 Modulation of competence for genetic transformation in Streptococcus pneumoniae. J. Gen. Microbiol. 133, (1959–67). (doi:10.1099/00221287-133-7-1959)

41. Wilson, G. A. & Bott, K. F. 1968 Nutritional factors influencing the development of competence in the Bacillus subtilis transformation system. J. Bacteriol. 95, 1439–49.

42. Palmen, R., Vosman, B., Buijsman, P., Breek, C. K. & Hellingwerf, K. J. 1993 Physiological characterization of natural transformation in Acinetobacter calcoaceticus. J. Gen. Microbiol. 139, (295–305). (doi:10.1099/00221287-139-2-295)

43. Kim, J.-S., Kim, J.-W. & Kathariou, S. 2008 Differential effects of temperature on natural transformation to erythromycin and nalidixic acid resistance in Campylobacter coli. Appl. Environ. Microbiol. 74, (6121–5). (doi:10.1128/AEM.01075-08)

44. Otto, S. P. 2009 The evolutionary enigma of sex. Am. Nat. 174 Suppl, S1–S14. (doi:10.1086/599084)

45. Hartfield, M. & Keightley, P. D. 2012 Current hypotheses for the evolution of sex and recombination. Integr. Zool. 7, (192–209). (doi:10.1111/j.1749-4877.2012.00284.x)

46. Vos, M. 2009 Why do bacteria engage in homologous recombination? Trends Microbiol. 17, (226–32). (doi:10.1016/j.tim.2009.03.001)

47. Altenberg, L. & Feldman, M. W. 1987 Selection, generalized transmission and the evolution of modifier genes. I. The reduction principle. Genetics 117, 559–72.

48. Kondrashov, A. S. 1984 Deleterious mutations as an evolutionary factor. 1. The advantage of recombination. Genet. Res. 44, 199–217.

49. Redfield, R. J. 1988 Evolution of bacterial transformation: is sex with dead cells ever better than no sex at all? Genetics 119, 213–221.

50. Redfield, R. J., Schrag, M. R. & Dean, A. M. 1997 The evolution of bacterial transformation: sex with poor relations. Genetics 146, 27–38.

51. de Visser, J. A. G. M. & Elena, S. F. 2007 The evolution of sex: empirical insights into the roles of epistasis and drift. Nat. Rev. Genet. 8, (139–49). (doi:10.1038/nrg1985)

52. Kouyos, R. D., Silander, O. K. & Bonhoeffer, S. 2007 Epistasis between deleterious mutations and the evolution of recombination. Trends Ecol. Evol. 22, (308–15). (doi:10.1016/j.tree.2007.02.014)

53. Hill, W. G. & Robertson, A. 1966 The effect of linkage on limits to artificial selection. Genet. Res. 8, 269–94.

54. Felsenstein, J. 1974 The evolutionary advantage of recombination. Genetics 78, 737–56.

55. Barton, N. H. & Otto, S. P. 2005 Evolution of recombination due to random drift]. Genetics 169, (2353–70). (doi:10.1534/genetics.104.032821)

56. Keightley, P. D. & Otto, S. P. 2006 Interference among deleterious mutations favours sex and recombination in finite populations. Nature 443, (89–92). (doi:10.1038/nature05049)

57. Fisher, R. A. 1930 The Genetical Theory Of Natural SelectionL: Fisher, R. AL: Free Download & StreamingL: Internet Archive. [cited 2016 Apr. 1].

58. Muller, H. J. 1932 Some Genetic Aspects of Sex. Am. Nat. 66, (118–138). (doi:10.1086/280418)

59. Muller, H. J. 1964 The relation of recombination to mutational advance. Mutat. Res. Mol. Mech. Mutagen. 1, (2–9). (doi:10.1016/0027-5107(64)90047-8)

60. Takeuchi, N., Kaneko, K. & Koonin, E. V 2014 Horizontal gene transfer can rescue prokaryotes from Muller’s ratchet: benefit of DNA from dead cells and population subdivision. G3 (Bethesda). 4, (325–39). (doi:10.1534/g3.113.009845)

61. Levin, B. R. & Cornejo, O. E. 2009 The population and evolutionary dynamics of homologous gene recombination in bacterial populations. PLoS Genet. 5, e1000601. (doi:10.1371/journal.pgen.1000601)

62. Moradigaravand, D. & Engelstädter, J. 2013 The evolution of natural competence: disentangling costs and benefits of sex in bacteria. Am. Nat. 182, (E112–26). (doi:10.1086/671909)

63. Cooper, T. F. 2007 Recombination speeds adaptation by reducing competition between beneficial mutations in populations of Escherichia coli. PLoS Biol. 5, e225. (doi:10.1371/journal.pbio.0050225)

64. McDonald, M. J., Rice, D. P. & Desai, M. M. 2016 Sex speeds adaptation by altering the dynamics of molecular evolution. Nature 531, (233–236). (doi:10.1038/nature17143)

65. Baltrus, D. A., Guillemin, K. & Phillips, P. C. 2008 Natural transformation increases the rate of adaptation in the human pathogen Helicobacter pylori. Evolution 62, (39–49). (doi:10.1111/j.1558-5646.2007.00271.x)

66. Bacher, J. M., Metzgar, D. & de Crécy-Lagard, V. 2006 Rapid evolution of diminished transformability in Acinetobacter baylyi. J. Bacteriol. 188, (8534–42). (doi:10.1128/JB.00846-06)

67. Utnes, A. L. G., S∅rum, V., Hülter, N., Primicerio, R., Hegstad, J., Kloos, J., Nielsen, K. M. & Johnsen, P. J. 2015 Growth phase-specific evolutionary benefits of natural transformation in Acinetobacter baylyi. ISME J. 9, (2221–31). (doi:10.1038/ismej.2015.35)

68. Engelmoer, D. J. p., Donaldson, I. & Rozen, D. E. 2013 Conservative sex and the benefits of transformation in Streptococcus pneumoniae. PLoS Pathog. 9, e1003758. (doi:10.1371/journal.ppat.1003758)

69. Redfield, R. J. 1988 Evolution of bacterial transformation: is sex with dead cells ever better than no sex at all? Genetics 119, 213–21.

70. Moradigaravand, D. & Engelstädter, J. 2014 The impact of natural transformation on adaptation in spatially structured bacterial populations. BMC Evol. Biol. 14, 141. (doi:10.1186/1471-2148-14-141)

71. Engelstädter, J. & Moradigaravand, D. 2014 Adaptation through genetic time travel? Fluctuating selection can drive the evolution of bacterial transformation. Proc. Biol. Sci. 281, 20132609. (doi:10.1098/rspb.2013.2609)

72. Croucher, N. J., Mostowy, R., Wymant, C., Turner, P., Bentley, S. D. & Fraser, C. 2016 Horizontal DNA Transfer Mechanisms of Bacteria as Weapons of Intragenomic Conflict. PLOS Biol. 14, e1002394. (doi:10.1371/journal.pbio.1002394)

73. Majewski, J. & Cohan, F. M. 1999 DNA sequence similarity requirements for interspecific recombination in Bacillus. Genetics 153, 1525–33.

74. Johnston, C., Martin, B., Granadel, C., Polard, P. & Claverys, J.-P. 2013 Programmed protection of foreign DNA from restriction allows pathogenicity island exchange during pneumococcal transformation. PLoS Pathog. 9, e1003178. (doi:10.1371/journal.ppat.1003178)

75. Croucher, N. J., Harris, S. R., Barquist, L., Parkhill, J. & Bentley, S. D. 2012 A high-resolution view of genome-wide pneumococcal transformation. PLoS Pathog. 8, e1002745. (doi:10.1371/journal.ppat.1002745)

76. Mell, J. C., Lee, J. Y., Firme, M., Sinha, S. & Redfield, R. J. 2014 Extensive cotransformation of natural variation into chromosomes of naturally competent Haemophilus influenzae. G3 (Bethesda). 4, (717–31). (doi:10.1534/g3.113.009597)

77. Fouts, D. E. 2006 Phage_Finder: automated identification and classification of prophage regions in complete bacterial genome sequences. Nucleic Acids Res. 34, (5839–51). (doi:10.1093/nar/gkl732)

78. West, S. A., Lively, C. M. & Read, A. F. 1999 A pluralist approach to sex and recombination. J. Evol. Biol. 12, (1003–1012). (doi:10.1046/j.1420-9101.1999.00119.x)

79. Charpentier, X., Polard, P. & Claverys, J.-P. 2012 Induction of competence for genetic transformation by antibiotics: convergent evolution of stress responses in distant bacterial species lacking SOS? Curr. Opin. Microbiol. 15, (570–6). (doi:10.1016/j.mib.2012.08.001)

80. Biswas, G. D., Thompson, S. A. & Sparling, P. F. 1989 Gene transfer in Neisseria gonorrhoeae. Clin. Microbiol. Rev. 2 Suppl, S24–8.

81. Smillie, C. S., Smith, M. B., Friedman, J., Cordero, O. X., David, L. A. & Alm, E. J. 2011 Ecology drives a global network of gene exchange connecting the human microbiome. Nature 480, (241–4). (doi:10.1038/nature10571)

82. Meibom, K. L., Blokesch, M., Dolganov, N. A., Wu, C.-Y. & Schoolnik, G. K. 2005 Chitin induces natural competence in Vibrio cholerae. Science 310, (1824–7). (doi:10.1126/science.1120096)

83. Suckow, G., Seitz, P. & Blokesch, M. 2011 Quorum sensing contributes to natural transformation of Vibrio cholerae in a species-specific manner. J. Bacteriol. 193, (4914–24). (doi:10.1128/JB.05396-11)

84. Yang, J., Evans, B. A. & Rozen, D. E. 2010 Signal diffusion and the mitigation of social exploitation in pneumococcal competence signalling. Proc. Biol. Sci. 277, (2991–9). (doi:10.1098/rspb.2010.0659)

85. Cornejo, O. E., McGee, L. & Rozen, D. E. 2010 Polymorphic competence peptides do not restrict recombination in Streptococcus pneumoniae. Mol. Biol. Evol. 27, (694–702). (doi:10.1093/molbev/msp287)

86. Stefanic, P. & Mandic-Mulec, I. 2009 Social interactions and distribution of Bacillus subtilis pherotypes at microscale. J. Bacteriol. 191, (1756–64). (doi:10.1128/JB.01290-08)

87. Iannelli, F., Oggioni, M. R. & Pozzi, G. 2005 Sensor domain of histidine kinase ComD confers competence pherotype specificity in Streptoccoccus pneumoniae. FEMS Microbiol. Lett. 252, (321–6). (doi:10.1016/j.femsle.2005.09.008)

88. Carrolo, M., Pinto, F. R., Melo-Cristino, J. & Ramirez, M. 2009 Pherotypes are driving genetic differentiation within Streptococcus pneumoniae. BMC Microbiol. 9, 191. (doi:10.1186/1471-2180-9-191)

89. Steinmoen, H., Teigen, A. & Havarstein, L. S. 2003 Competence-Induced Cells of Streptococcus pneumoniae Lyse Competence-Deficient Cells of the Same Strain during Cocultivation. J. Bacteriol. 185, (7176–7183). (doi:10.1128/JB.185.24.7176-7183.2003)

90. Guiral, S., Mitchell, T. J., Martin, B. & Claverys, J.-P. 2005 Competence-programmed predation of noncompetent cells in the human pathogen Streptococcus pneumoniae: genetic requirements. Proc. Natl. Acad. Sci. U. S. A. 102, (8710–5). (doi:10.1073/pnas.0500879102)

91. Claverys, J.-P., Martin, B. & Håvarstein, L. S. 2007 Competence-induced fratricide in streptococci. Mol. Microbiol. 64, (1423–33). (doi:10.1111/j.1365-2958.2007.05757.x)

92. Kjos, M., Miller, E., Slager, J., Lake, F. B., Gericke, O., Roberts, I. S., Rozen, D. E. & Veening, J.-W. 2016 Expression of Streptococcus pneumoniae Bacteriocins Is Induced by Antibiotics via Regulatory Interplay with the Competence System. PLoS Pathog. 12, e1005422. (doi:10.1371/journal.ppat.1005422)

93. Wholey, W.-Y., Kochan, T. J., Storck, D. N. & Dawid, S. 2016 Coordinated Bacteriocin Expression and Competence in Streptococcus pneumoniae Contributes to Genetic Adaptation through Neighbor Predation. PLoS Pathog. 12, e1005413. (doi:10.1371/journal.ppat.1005413)

94. Cianfanelli, F. R., Monlezun, L. & Coulthurst, S. J. 2015 Aim, Load, Fire: The Type VI Secretion System, a Bacterial Nanoweapon. Trends Microbiol. 24, (51–62). (doi:10.1016/j.tim.2015.10.005)

95. Borgeaud, S., Metzger, L. C., Scrignari, T. & Blokesch, M. 2015 The type VI secretion system of Vibrio cholerae fosters horizontal gene transfer. Science 347, (63–7). (doi:10.1126/science.1260064)

96. Duffin, P. M. & Seifert, H. S. 2010 DNA uptake sequence-mediated enhancement of transformation in Neisseria gonorrhoeae is strain dependent. J. Bacteriol. 192, (4436–44). (doi:10.1128/JB.00442-10)

97. Cehovin, A. et al. 2013 Specific DNA recognition mediated by a type IV pilin. Proc. Natl. Acad. Sci. U. S. A. 110, (3065–70). (doi:10.1073/pnas.1218832110)

98. Goodman, S. D. & Scocca, J. J. 1988 Identification and arrangement of the DNA sequence recognized in specific transformation of Neisseria gonorrhoeae. Proc. Natl. Acad. Sci. U. S. A. 85, 6982–6.

99. Frye, S. A., Nilsen, M., T∅njum, T. & Ambur, O. H. 2013 Dialects of the DNA uptake sequence in Neisseriaceae. PLoS Genet. 9, e1003458. (doi:10.1371/journal.pgen.1003458)

100. Redfield, R. J., Findlay, W. A., Bossé, J., Kroll, J. S., Cameron, A. D. S. & Nash, J. H. 2006 Evolution of competence and DNA uptake specificity in the Pasteurellaceae. BMC Evol. Biol. 6, 82. (doi:10.1186/1471-2148-6-82)

101. Treangen, T. J., Ambur, O. H., Tonjum, T. & Rocha, E. P. C. 2008 The impact of the neisserial DNA uptake sequences on genome evolution and stability. Genome Biol. 9, R60. (doi:10.1186/gb-2008-9-3-r60)

102. Ambur, O. H., Frye, S. A., Nilsen, M., Hovland, E. & T∅njum, T. 2012 Restriction and sequence alterations affect DNA uptake sequence-dependent transformation in Neisseria meningitidis. PLoS One 7, e39742. (doi:10.1371/journal.pone.0039742)

103. Ambur, O. H., Frye, S. A. & T∅njum, T. 2007 New functional identity for the DNA uptake sequence in transformation and its presence in transcriptional terminators. J. Bacteriol. 189, (2077–85). (doi:10.1128/JB.01408-06)

104. Mell, J. C., Hall, I. M. & Redfield, R. J. 2012 Defining the DNA uptake specificity of naturally competent Haemophilus influenzae cells. Nucleic Acids Res. 40, (8536–49). (doi:10.1093/nar/gks640)

105. Davidsen, T., R∅dland, E. A., Lagesen, K., Seeberg, E., Rognes, T. & T∅njum, T. 2004 Biased distribution of DNA uptake sequences towards genome maintenance genes. Nucleic Acids Res. 32, (1050–8). (doi:10.1093/nar/gkh255)

106. Oliveira, P. H., Touchon, M. & Rocha, E. P. C. 2014 The interplay of restriction-modification systems with mobile genetic elements and their prokaryotic hosts. Nucleic Acids Res. 42, (10618–31). (doi:10.1093/nar/gku734)

107. Roberts, R. J., Vincze, T., Posfai, J. & Macelis, D. 2015 REBASE–a database for DNA restriction and modification: enzymes, genes and genomes. Nucleic Acids Res. 43, (D298–9). (doi:10.1093/nar/gku 1046)

108. Berndt, C., Meier, P. & Wackernagel, W. 2003 DNA restriction is a barrier to natural transformation in Pseudomonas stutzeri JM300. Microbiology 149, (895–901). (doi:10.1099/mic.0.26033-0)

109. Morikawa, K., Takemura, A. J., Inose, Y., Tsai, M., Nguyen Thi, L. T., Ohta, T. & Msadek, T. 2012 Expression of a cryptic secondary sigma factor gene unveils natural competence for DNA transformation in Staphylococcus aureus. PLoS Pathog. 8, e1003003. (doi:10.1371/journal.ppat.1003003)

110. Fagerlund, A., Granum, P. E. & Håvarstein, L. S. 2014 Staphylococcus aureus competence genes: mapping of the SigH, ComK1 and ComK2 regulons by transcriptome sequencing. Mol. Microbiol. 94, (557–79). (doi:10.1111/mmi.12767)

111. Chung, D., Farkas, J. & Westpheling, J. 2013 Overcoming restriction as a barrier to DNA transformation in Caldicellulosiruptor species results in efficient marker replacement. Biotechnol. Biofuels 6, 82. (doi:10.1186/1754-6834-6-82)

112. Ando, T., Xu, Q., Torres, M., Kusugami, K., Israel, D. A. & Blaser, M. J. 2000 Restriction-modification system differences in Helicobacter pylori are a barrier to interstrain plasmid transfer. Mol. Microbiol. 37, 1052–65.

113. Jeltsch, A. 2003 Maintenance of species identity and controlling speciation of bacteria: a new function for restriction/modification systems? Gene 317, 13–6.

114. Budroni, S. et al. 2011 Neisseria meningitidis is structured in clades associated with restriction modification systems that modulate homologous recombination. Proc. Natl. Acad. Sci. U. S. A. 108, (4494–9). (doi:10.1073/pnas.1019751108)

115. Godde, J. S. & Bickerton, A. 2006 The repetitive DNA elements called CRISPRs and their associated genes: evidence of horizontal transfer among prokaryotes. J. Mol. Evol. 62, 718–29. (doi:10.1007/s00239-005-0223-z)

116. Bikard, D., Hatoum-Aslan, A., Mucida, D. & Marraffini, L. A. 2012 CRISPR interference can prevent natural transformation and virulence acquisition during in vivo bacterial infection. Cell Host Microbe 12, (177–86). (doi:10.1016/j.chom.2012.06.003)

117. Jorth, P. & Whiteley, M. 2012 An evolutionary link between natural transformation and CRISPR adaptive immunity. MBio 3. (doi:10.1128/mBio.00309-12)

118. Gophna, U., Kristensen, D. M., Wolf, Y. I., Popa, O., Drevet, C. & Koonin, E. V 2015 No evidence of inhibition of horizontal gene transfer by CRISPR-Cas on evolutionary timescales. ISME J. 9, (2021–7). (doi:10.1038/ismej.2015.20)

119. Kowalczykowski, S. C., Dixon, D. A., Eggleston, A. K., Lauder, S. D. & Rehrauer, W. M. 1994 Biochemistry of homologous recombination in Escherichia coli. Microbiol. Rev. 58, 401–65.

120. Mortier-Barrière, I. et al. 2007 A key presynaptic role in transformation for a widespread bacterial protein: DprA conveys incoming ssDNA to RecA. Cell 130, (824–36). (doi:10.1016/j.cell.2007.07.038)

121. Yadav, T., Carrasco, B., Serrano, E. & Alonso, J. C. 2014 Roles of Bacillus subtilis DprA and SsbA in RecA-mediated genetic recombination. J. Biol. Chem. 289, (27640–52). (doi:10.1074/jbc.M114.577924)

122. Carrasco, B., Yadav, T., Serrano, E. & Alonso, J. C. 2015 Bacillus subtilis RecO and SsbA are crucial for RecA-mediated recombinational DNA repair. Nucleic Acids Res. 43, (5984–97). (doi:10.1093/nar/gkv545)

123. Sagi, D., Tlusty, T. & Stavans, J. 2006 High fidelity of RecA-catalyzed recombination: a watchdog of genetic diversity. Nucleic Acids Res. 34, (5021–31). (doi:10.1093/nar/gkl586)

124. Honigberg, S. M., Rao, B. J. & Radding, C. M. 1986 Ability of RecA protein to promote a search for rare sequences in duplex DNA. Proc. Natl. Acad. Sci. U. S. A. 83, 9586–90.

125. Roberts, M. S. & Cohan, F. M. 1993 The effect of DNA sequence divergence on sexual isolation in Bacillus. Genetics 134, 401–8.

126. Zawadzki, P., Roberts, M. S. & Cohan, F. M. 1995 The log-linear relationship between sexual isolation and sequence divergence in Bacillus transformation is robust. Genetics 140, 917–32.

127. Vulić, M., Dionisio, F., Taddei, F. & Radman, M. 1997 Molecular keys to speciation: DNA polymorphism and the control of genetic exchange in enterobacteria. Proc. Natl. Acad. Sci. U. S. A. 94, 9763–7.

128. Eppley, J. M., Tyson, G. W., Getz, W. M. & Banfield, J. F. 2007 Genetic exchange across a species boundary in the archaeal genus ferroplasma. Genetics 177, (407–16). (doi:10.1534/genetics.107.072892)

129. Ray, J. L., Harms, K., Wikmark, O.-G., Starikova, I., Johnsen, P. J. & Nielsen, K. M. 2009 Sexual isolation in Acinetobacter baylyi is locus-specific and varies 10,000-fold over the genome. Genetics 182, (1165–81). (doi:10.1534/genetics.109.103127)

130. Majewski, J., Zawadzki, P., Pickerill, P., Cohan, F. M. & Dowson, C. G. 2000 Barriers to Genetic Exchange between Bacterial Species: Streptococcus pneumoniae Transformation. J. Bacteriol. 182, (1016–1023). (doi:10.1128/JB.182.4.1016-1023.2000)

131. Frost, L. S., Leplae, R., Summers, A. O. & Toussaint, A. 2005 Mobile genetic elements: the agents of open source evolution. Nat. Rev. Microbiol. 3, (722–32). (doi:10.1038/nrmicro1235)

132. Ali, S. S., Soo, J., Rao, C., Leung, A. S., Ngai, D. H.-M., Ensminger, A. W. & Navarre, W. W. 2014 Silencing by H-NS potentiated the evolution of Salmonella. PLoS Pathog. 10, e1004500. (doi:10.1371/journal.ppat.1004500)

133. Navarre, W. W., Porwollik, S., Wang, Y., McClelland, M., Rosen, H., Libby, S. J. & Fang, F. C. 2006 Selective silencing of foreign DNA with low GC content by the H-NS protein in Salmonella. Science 313, (236–8). (doi:10.1126/science.1128794)

134. Baltrus, D. A. 2013 Exploring the costs of horizontal gene transfer. Trends Ecol. Evol. 28, (489–95). (doi:10.1016/j.tree.2013.04.002)

135. Knöppel, A., Lind, P. A., Lustig, U., Näsvall, J. & Andersson, D. I. 2014 Minor fitness costs in an experimental model of horizontal gene transfer in bacteria. Mol. Biol. Evol. 31, (1220–7). (doi:10.1093/molbev/msu076)

136. Bershtein, S., Serohijos, A. W. R., Bhattacharyya, S., Manhart, M., Choi, J.-M., Mu, W., Zhou, J. & Shakhnovich, E. I. 2015 Protein Homeostasis Imposes a Barrier on Functional Integration of Horizontally Transferred Genes in Bacteria.PLoS Genet. 11, e1005612. (doi:10.1371/journal.pgen.1005612)

